# Modelling mechanisms and treatment of cholangiopathies with a bile duct on a chip

**DOI:** 10.64898/2026.08.19.745387

**Authors:** Henry William Hoyle, Anna Katharina Frank, Enya Amundsen-Isaksen, Sarah Peisl, Oline Øie Hovland, Jeremy Yeoh, Mughilan Selvarajah, Aleksandra Aizenshtadt, Kayoko Hirayama-Shoji, Fotios Sampaziotis, Tom Hemming Karlsen, Mathias Busek, Stefan Krauss, Espen Melum

## Abstract

**Background and aims:** Model systems for bile duct disorders are needed for testing therapeutic interventions. Current models have poor human relevance or limited potential for recreating the complex bile duct microenvironment at scale. We aimed to generate a humanized microphysiological system to model and treat cholangiopathies.

**Methods:** An *in vitro* bile duct was created using 3D printed microfluidic chips containing a collagen-embedded canal seeded with patient-derived primary human cholangiocytes. Barrier permeability and compound transport across the epithelium was measured, and disruption of the barrier was performed with lipopolysaccharide treatment. The duct was challenged with the known hepatotoxicant Chlorpromazine. Biliatresone was used to model biliary-atresia and treated using N-acetyl-L-cysteine.

**Results:** Cholangiocytes in the bile duct chip established a tight, polarized epithelial barrier. Verapamil and Linerixibat inhibited transport of rhodamine 123 and cholyl-lys-fluorescein respectively with 66 % (p = 0.0004) and 57 % (p = 0.03) reduction. 10 µg/mL lipopolysaccharide led to a loss of epithelial barrier integrity, measured by an increase of over 1000 % in leakage of both 3 kDa (p = 0.0002) and 10 kDa dextran (p = 0.0001) along with upregulation of cytokines. Chlorpromazine displayed dose-dependent toxicity with EC50 values of 84, 140 and 96 µM for three patient lines. Biliatresone induced a dose-dependent abnormal phenotype with loss of viability. The induced phenotype could be treated with N-acetyl-L-cysteine, improving viability from 23 % to 59 % (p < 0.0001) with treatment of 2 µg/mL Biliatresone.

**Conclusions:** Our novel platform allows complex studies of bile duct biology, testing of off-target effects from drugs and treatment of a disease phenotype.

## Introduction

Bile duct diseases constitute a number of debilitating conditions such as primary sclerosing cholangitis (PSC), primary biliary cholangitis (PBC) in adults and Alagille syndrome (AS) and biliary atresia (BA) in children^1^. The diseases often involve genetic, environmental and idiopathic factors, however the exact etiology is unclear in most of the bile duct diseases. Clinical presentations include inflammation, strictures, and cholestasis, affecting the normal flow of bile leading to altered composition and leakage into the liver. They place a heavy burden on patients due to poor prognosis, symptoms such as pruritus and jaundice, and increased risk of cirrhosis, liver failure and cholangiocarcinoma^2^. Limited treatment options and rising prevalence highlight the need for improved understanding and more effective treatment modalities^3^. These challenges can be hindered by a lack of high-quality preclinical models, where mouse models are considered the gold standard yet suffer from shortcomings such as incomplete recapitulation of disease characteristics and reduced relevance to human disease^4^.

A range of *in vitro* models of the bile duct have been developed to allow studies to be performed using human material. The most basic studies utilize monolayer cultures of immortalized cholangiocyte cell lines^5^ however these are not tailored to specific patients and diseases. Primary cholangiocytes on the other hand have limited value in monolayers due to a short functional lifespan and loss of cellular polarization^6^. Cholangiocyte organoid cultures, where spheroids of cells are grown within an extracellular matrix gel, are increasingly being used due to the ability to maintain primary cells in a differentiated state for prolonged periods. These can utilize cell sources such as stem cells^7^ and liver, extrahepatic bile duct or gallbladder biopsies^8–10^. Despite their advantages, cholangiocyte organoids still lack a structure resembling the bile duct and their interior, representing the inside of the *in vivo* bile duct, cannot easily be reached.

Increased complexity and physiological relevance can be achieved with organ-on-a-chip (OoC) technology, allowing cell growth in a defined geometry with options to include mechanical stimuli such as fluid flow^11–13^. The spatial control of cellular elements can be achieved by modern fabrication techniques where a range of wells, compartments and interconnecting conduits can be produced^14^. OoC approaches with primary cholangiocytes in a tubular structure have demonstrated close resemblance to the *in vivo* bile duct structure^15–17^ however suffer from drawbacks such as limited scalability due to the use of soft lithography for fabrication^18^ and high levels of drug absorption with hydrophobic materials such as polydimethylsiloxane (PDMS)^19^.

We herein aimed to leverage technological advances and establish a bile duct on a chip which exhibits high physiological relevance and scalability alongside technical simplicity. The primary outcome is to facilitate more efficient development of novel treatments for bile duct diseases however this could also have future use in personalized medicine to address patient heterogeneity in the cholangiopathies by identifying optimal treatments for individual patients. Alongside generating robust data about cholangiocyte function and disease, we also demonstrate the utility for clinically relevant drug studies across a range of pathways such as transport modulation and cytotoxicity, as well as challenge through the addition of lipopolysaccharide (LPS). Finally, we model and treat BA in the chip using the compound Biliatresone, highlighting the potential for generating robust disease models for testing novel treatment.

## Materials and Methods

### Patient material and organoid culture

Cholangiocytes were obtained from the bile ducts of patients undergoing ERCP and cultured as organoids as previously described^9^. The patients were recruited at the Section of Gastroenterology at the Department of Transplantation Medicine, Oslo University Hospital Rikshospitalet. Written informed consent was obtained prior to ERCP and the collection and use of material by the Norwegian PSC research center biobanks has been approval by the Regional Ethical Committee (REK 15368, 599798 and 18221). Patient inclusion in the study was performed independent of gender (**Table S1**). All organoids were derived from patients diagnosed with PSC due to ease of access to material and previous data showing minimal difference between healthy and diseased cholangiocytes when cultured as organoids^20^. All samples were routinely tested for mycoplasma (MycoAlert PLUS, Lonza, Basel, Switzerland).

### Chip fabrication and culture

The design for the chip was created using Solidworks 2020 (Dassault Systèmes, Vélizy-Villacoublay, France). Files were arranged for printing using PreForm 3.31.0 (FormLabs, Somerville, Massachusetts, USA). Chips were printed using a Form4 3D printer (FormLabs) using High Temperature Resin V2 (FormLabs). After printing chips were washed twice for 10 minutes each in isopropanol using a Form Wash V2 (FormLabs) with drying using compressed air between washes and then cured using a Form Cure V2 (FormLabs) for 20 minutes at 80°C. A 13.5 mm x 53 mm coverslip was laser cut from 100 µm thick Zeonor® film with a glass transition temperature of 163 °C (ZEON Corporation, Tokyo, Japan) and adhered to the cured chips using High Temperature Resin V2. The resin was cured under 405nm light for 1 minute. Chips were washed for 1 minute in fresh isopropanol and dried using compressed air before being cured in the Form Cure V2 for 20 minutes at 80°C. Finally, chips were autoclaved for 20 minutes at 121°C prior to use.

### Formation of a collagen gel in the chip

Autoclaved chips were removed from autoclave pouches in a sterile cell culture cabinet. 70 µL of 0.01 % Poly-L-lysine (PLL, Sigma-Aldrich) solution was added to the basal wells of each chip and incubated for 2 hours at room temperature. PLL was aspirated and chips washed with sterile distilled water before being left to air dry. Alignment inserts were placed into access ports on the left hand side of chips and autoclaved 200 µm diameter glass capillaries (Capillary tube supplies Ltd, Cornwall, UK) inserted into each channel. Collagen solution was made up on ice in 10X Phosphate buffered saline (PBS, Gibco) with 4 mg/ml CellAdhere™ Type I bovine collagen (STEMCELL Technologies, Vancouver, Canada), 1 mg/mL high concentration rat tail collagen I (Corning) and 16.7 mM sodium hydroxide. The ice-cold collagen solution was degassed in a vacuum chamber prior to use. 110 µL of collagen solution was added to the central well of each channel and incubated at 37 °C for 2 hours to polymerize the collagen. Chips were washed with PBS and the glass capillaries removed to expose the channels. Thermoplastic mini luer end caps (MicroFluidic Chip Shop, Jena, Germany) were fitted to the chips and the PBS replaced with Williams medium E prior to storage at 37 °C (< 3 days) or RT (< 1 week).

### Culture of cholangiocytes in the bile duct chip

Organoids were washed with cell recovery solution (Corning) and transferred to a centrifuge tube. The tubes were incubated for 30 minutes on ice with resuspension every 10 minutes. During this time the prepared chips were washed once with wash medium.

After 30 minutes the organoid suspension was centrifuged at 482 g for 4 minutes. Supernatant was removed and organoids resuspended in 10 mL of wash medium. A second 482 g, 4-minute centrifugation was performed, and organoids were resuspended in 2 mL of prewarmed 0.25 % Trypsin-EDTA (Gibco). The suspension was incubated in a 37 °C water bath for 6 minutes, with resuspension with a pipette every 1.5 minutes. After this time, 8 mL of wash medium supplemented with 1 % fetal bovine serum (FBS, Gibco) was added and tubes were centrifuged for 8 minutes at 544 g. Supernatant was removed and cells were washed through a 70 µm cell strainer in 15 mL of wash medium. Cells were centrifuged at 544 g for 8 minutes and finally resuspended in growth medium with 10µM Y-27632. A cell count was performed with an EVE automated cell counter (NanoEntek, Seoul, South Korea) and cells were diluted to a concentration of 500,000 cells per mL. Medium was aspirated from the chips and 150 µL of cell suspension was added to each apical well, resulting in 150,000 cells seeded per channel. Chips were incubated at 37 °C for 2 hours and then media changed to remove excess cells. The following day, medium was aspirated, fresh growth medium was added and the chip was placed on a RoboRocker rocking platform (Next Advance, Troy, NY, USA) set to rock at a speed of 0.1 degrees per second. Subsequent media changes were performed every two days with growth medium.

All experiments were performed with chips at day 10. For full details of analytical techniques performed see supplementary materials and methods.

### Chip challenges

For LPS and CPZ challenge, fresh medium containing the compounds was added to the apical wells and medium without the compounds was added to the basal wells. Chips were incubated for 24 hours before performing further analysis. For BSO challenge, fresh medium containing BSO was added to all wells and chips were incubated for 48 hours before performing further analysis. See supplementary materials and methods for further details.

### Statistics

All experiments were performed with cholangiocytes from 3 different PSC patients, with 3-4 channels for each condition unless otherwise stated. Data is presented as mean ± SEM unless otherwise stated. Statistics were calculated using GraphPad Prism version 10.4.1 (GraphPad, San Diego, CA, USA). Comparisons between two groups were performed with Welch’s t-test while comparisons between multiple groups were performed using one-way ANOVA with Dunnett’s test for post-hoc comparison of treatment to control groups. Comparisons within grouped analyses were performed with two-way ANOVA using Šídák’s multiple comparisons test for post hoc comparisons between group means.

## Results

### Formation of a biliary epithelium in the microfluidic chip

To create an *in vitro* microfluidic model of the bile duct we employed a chip design allowing culture of cholangiocytes and circulation of fluids with high scalability, compatibility with standard analytical techniques and biocompatibility (**Figure 1A**). Using a stereolithography 3D printer we could use a complex design while at the same time having improved drug absorption properties compared to PDMS with for example a reduced absorption of hydrophobic compounds such as deoxycholic acid (**Figure S1A**). This production method was also highly scalable with 11 minutes production time per chip (2 minutes hands-on) when making batches of 24 (**Figure S1B**). After initial testing of different designs, we arrived at a design that optimized collagen stability though anchor points such as a flanged outlet for the duct channel and grooves set into the side walls (**Figure 1A, B, S1C**). To sterilize and increase biocompatibility the chips were autoclaved, demonstrating no loss in viability when cholangiocytes were grown as organoids in the presence of the plastic compared to substantial loss without autoclaving (**Figure S2A-C**). A collagen gel containing a straight channel was then created, with a diameter of 200 µm balancing the number of cells required to seed with physiologically relevant level of flow in range of ± 4 mm/s when rocked at 0.1 degrees/second (**Figure 1C, S2D**).

**Fig. 1.**
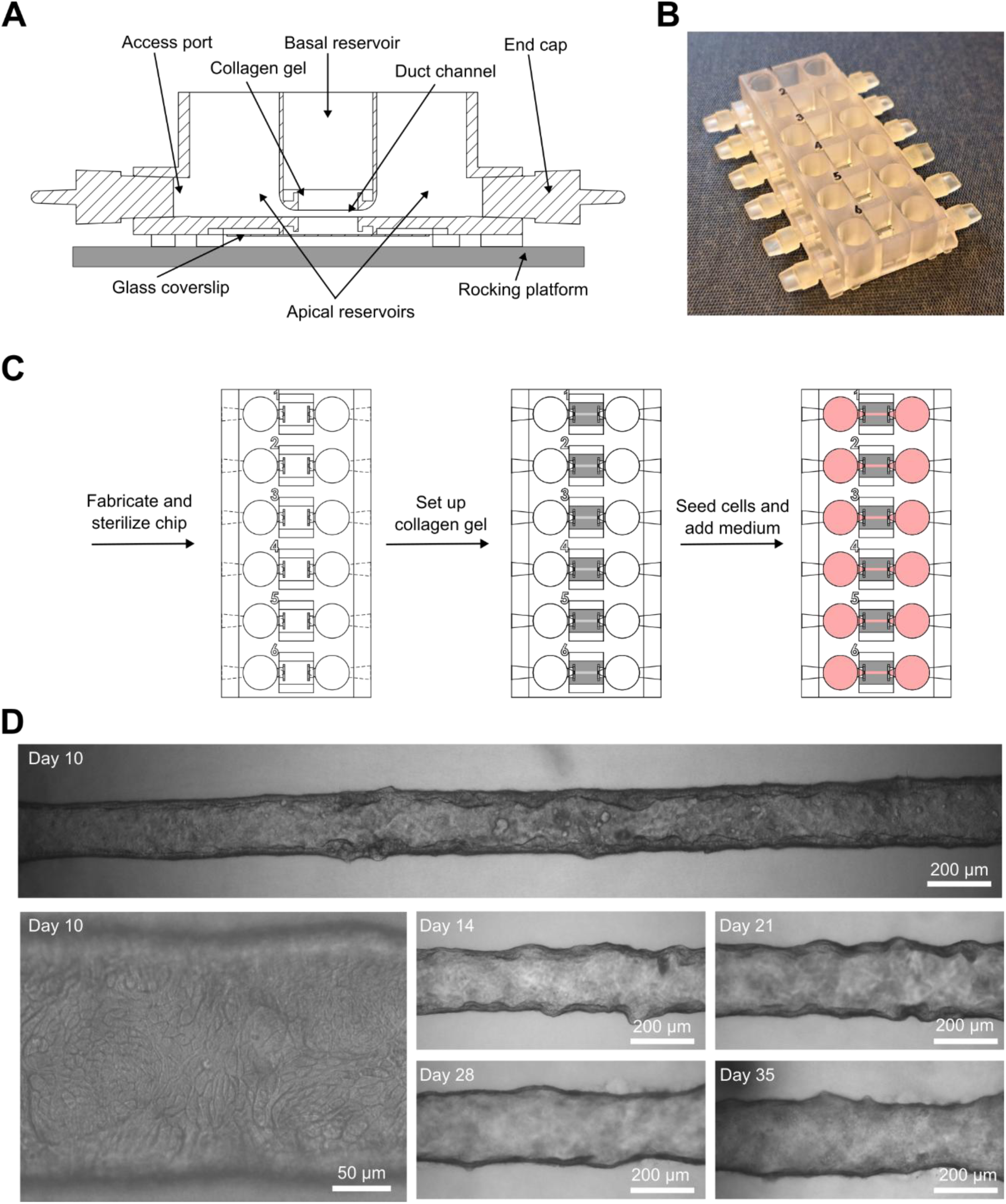
Formation of the bile duct on a chip. (A) Cross-section schematic of the bile duct chip. (B) Photo of the bile duct chip. (C) Schematic protocol for creation of the collagen gel within the chip. (D) Channels in the bile duct chip at days 10, 14, 21, 28 and 35. Images are representative data from one of three tested cholangiocyte patient lines (n=3-4). ERCP, Endoscopic retrograde cholangiopancreatography.

For establishment of the biliary epithelium inside the chip we used our previously developed pipeline for obtaining patient-derived primary cholangiocytes from ERCP brushes (**Figure S2E**) and cultured them as organoids (**Figure S2F**)^9^. Single cell suspensions of cholangiocyte organoids were seeded in the apical wells and after 7 days formed a confluent monolayered epithelium and reached a mature cholangiocyte morphology after 8-10 days and could be cultured for at least a further 3 weeks (**Figure 1D**). The mature cells in the chip exhibited an enhanced large cholangiocyte-like columnar phenotype compared to cholangiocyte organoids, measuring 5-10 µm in diameter and 15-20 µm in height with a mean width:height ratio of 0.4 in the chip compared to 1.0 of cholangiocytes in organoids (p < 0.0001) (**Figure S3A**)^21^.

Cholangiocytes cultured in the chip retained their cholangiocyte identity as evidenced by expression of cytokeratin 7, cytokeratin 19 and epithelial cell adhesion molecule (EPCAM) in the cells throughout the channels (**Figure 2A**). A sheet of filamentous actin formed localized towards the apical cell membranes and supported proper cholangiocyte polarization (**Figure 2A**). The cells in the channel also produced their own basement membrane around the outside of the channel, as evidenced by expression of collagen IV and laminin in this region (**Figure S3B**).

**Fig. 2.**
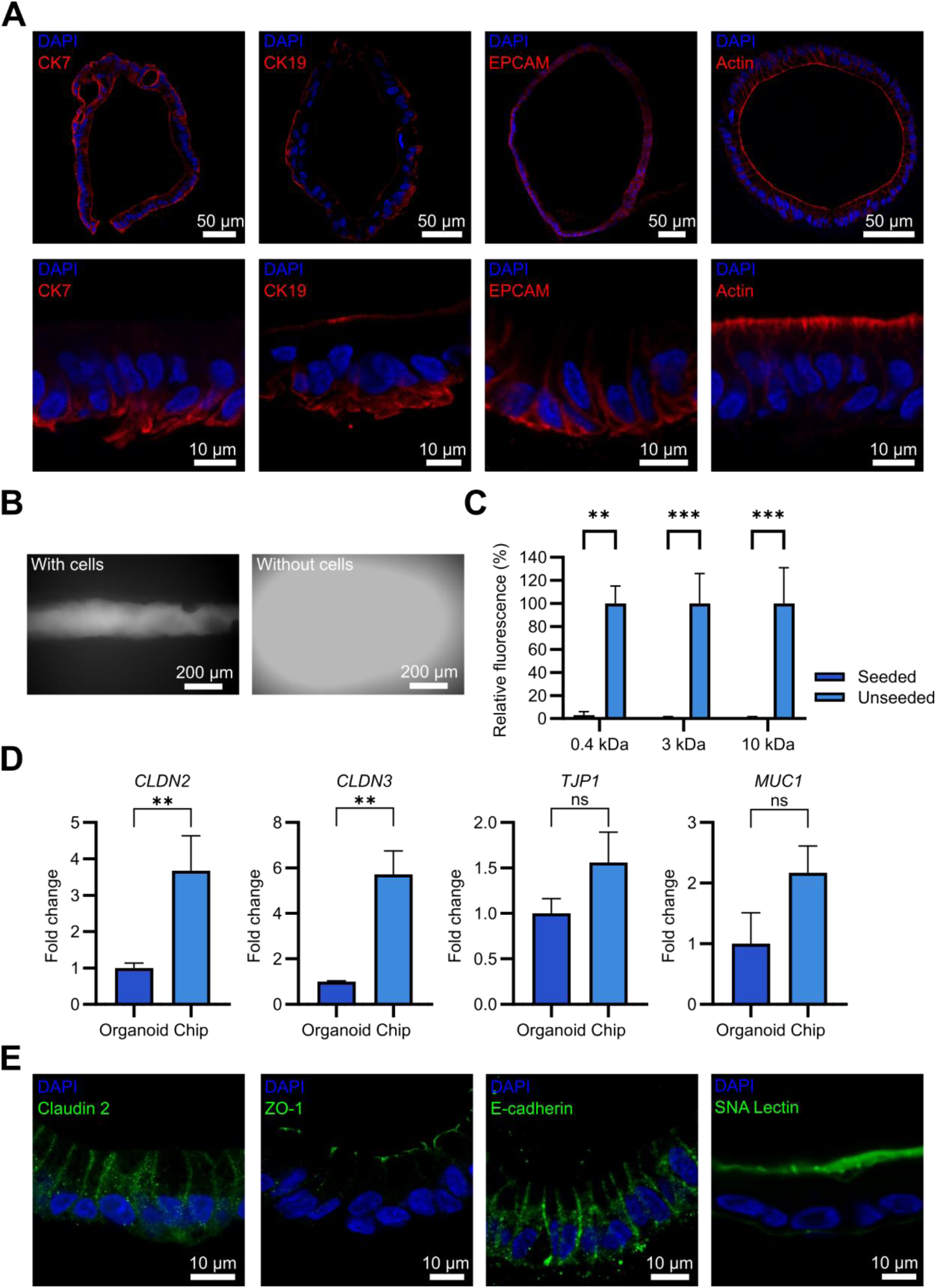
Basic characterization of the bile ducts generated *in vitro*. (A) Immunostaining for cholangiocyte identity proteins and actin. (B) Confocal microscope images of 10 kDa fluorescent dextran in channels with and without cells 5 minutes after addition. (C) Fluorescence in basal wells of different sized fluorescent compounds in channels with and without cholangiocytes after 1 hour of apical exposure (Mean ± SEM). (D) Fold change of RNA of epithelial barrier components in cholangiocytes grown as organoids and in the bile duct chip measured with RT-qPCR (Geometric mean ± SEM). (E) Immunostaining of proteins found in cell-cell junctions stained. SNA Lectin staining demonstrates the presence of mucus. All images and graphs are representative data from one of three tested cholangiocyte patient lines (n = 3-4). Statistical analyses were calculated using two-way ANOVA (C) or Welch’s t test (D). ns = p ≥ 0.05, ∗p < 0.05, ∗∗p < 0.01, and ∗∗∗p < 0.001, ∗∗∗∗p < 0.0001. CK7, Cytokeratin 7; CK19, Cytokeratin 19; EPCAM, Epithelial cell adhesion molecule; SNA, Sambucus Nigra; ZO-1, Zonula occludens-1.

### Epithelial barrier properties

After having demonstrated formation of a confluent polarized cholangiocyte channel in the chip, we next assessed the integrity of the epithelial barrier as this is important *in vivo* to prevent bile leakage into surrounding tissue. By using 0.4 kDa Lucifer yellow (LY), 3 kDa and 10 kDa fluorescent dextran molecules, the barrier was proven to be robust with minimal leakage when cholangiocytes were present in the channel compared to channels without cholangiocytes (3.1 %, 0.8 % and 1.2 % respectively, p = 0.001, 0.0009 and 0.0009) ruling out that diffusion limitations in the collagen gel could explain the barrier function (**Figure 2B, C**). Tight junctions are important in maintaining the epithelial barrier. Some genes related to barrier components in the bile duct on a chip have a higher expression than in comparable cholangiocyte organoids, with a change of 3.7-fold and 5.7-fold for *CLDN2* (p = 0.005) and *CLDN3* (p = 0.002) respectively, while *TJP1* is maintained at a similar level (**Figure 2D**). At the protein level the tight junction proteins claudin 2 and zonula occludens-1 (ZO-1) were clearly expressed at the cell membranes between adjacent cells, along with the adherens junction component E-Cadherin (**Figure 2E)**. The mucus layer is a key factor supporting barrier properties of the healthy bile duct^22^ and in the chip we observed *MUC1* expression at comparable levels to organoid cultures (**Figure 2D**) along with clear extracellular staining with SNA lectin, which binds to sialic acid, along the apical surface of the cholangiocytes (**Figure 2E).**

### Directional transport across the epithelium

We next assessed compound transport across the biliary epithelium, a key function for maintaining bile homeostasis. Several of the key transporters involved in these processes were upregulated in the bile duct chip compared to cholangiocyte organoids with significant fold change increases observed in *AE2* (3.6-fold, p = 0.001), *AQP1* (4.2-fold, p = 0.003), *ASBT* (8.2-fold, p = 0.002), *CFTR* (4.8-fold, p = 0.01) and *MDR1* (3.9-fold, p = 0.003) (**Figure 3A**).

**Fig. 3.**
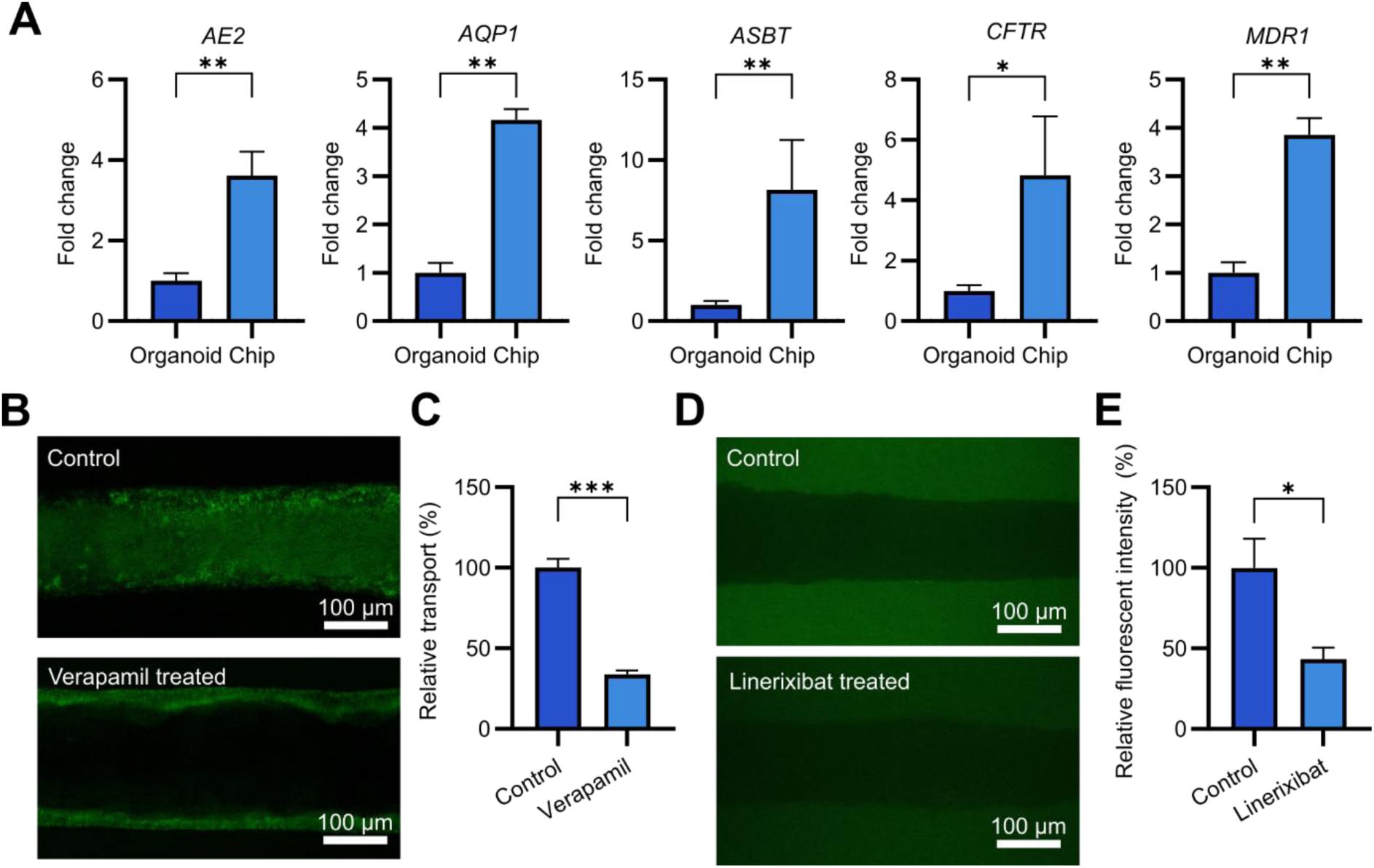
Demonstration of transporter expression and function in the artificial bile ducts. (A) Fold change of RNA for key transporters in cholangiocytes grown as organoids and in the bile duct chip measured with qPCR (Geometric mean ± SEM) (B) Confocal microscope images of channels in the bile duct chip 30 minutes after addition of rhodamine 123 to the basal wells with and without 20 µM Verapamil treatment. (C) Apical fluorescence relative to the control 30 minutes after addition of rhodamine 123 to the basal wells of the bile duct chip (Mean ± SEM). (D) Confocal microscope images of channels in the bile duct chip 30 minutes after addition of cholyl-lys-fluorescein to the apical wells with and without 50 µM Linerixibat treatment. (E) Basal fluorescence relative to the control 30 minutes after addition of cholyl-lys-fluorescein to the apical wells of the bile duct chip. (Mean ± SEM). All images and graphs are representative data from one of three tested cholangiocyte patient lines (n = 4). Statistical analyses were calculated using Welch’s t test. ns ≥ 0.05, ∗p < 0.05, ∗∗p < 0.01, and ∗∗∗p < 0.001, ∗∗∗∗p < 0.0001. AE2, anion exchanger 2; CFTR, cystic fibrosis transmembrane receptor.

The functionality of the membrane bound transporters was evaluated alongside inhibition with drugs, which is a relevant mechanism for potential future therapeutic treatments. Rhodamine 123, which is transported across the apical epithelium via MDR1, accumulated in the apical medium after addition to the basal wells (**Figure 3B, C**). Inhibition of MDR1 with the known inhibitor Verapamil, led to a 66 % reduction in the transport of rhodamine 123 into the apical wells (p = 0.0004) with a subsequent buildup within the cells.

Addition of CLF, a fluorescent-tagged bile acid, to the apical wells, led to build-up in the basal collagen. Inhibition of the key transporter ASBT using the drug Linerixibat, which is currently investigated for treatment of bile duct disorders in humans^23^ leads to a 57 % reduction in CLF transport out of the bile duct (p = 0.03) (**Figure 3D, E**).

### Challenge with a bacterial product increases barrier permeability at high doses

Our next aim was to demonstrate the response of the model to a range of physiological and disease relevant insults. We first investigated the impact of apical exposure to pathogen-associated molecular patterns (PAMPs). LPS, a commonly used cell wall component of gram-negative bacteria previously shown to cause damage to the bile duct barrier^24^, was used for this challenge. Addition of LPS to the bile ducts resulted in no major changes to the gross morphology of the channels after 24 hours (**Figure 4A**). Impacts on barrier integrity were measured with fluorescent dextran and LY. Leakage of LY through the biliary epithelium after 24 hours of LPS treatment was stable across all conditions. Both sizes of dextran molecules show no significant change in permeability after treatment with LPS at doses of 0.1 and 1 µg/mL, however a dose of 10 µg/mL led to a large increase in basal fluorescence for both 3 kDa dextran (1800 %, p = 0.0002) and 10 kDa dextran (1200 %, p = 0.0001) (**Figure 4B, C**). Upregulation of cytokines in the cholangiocytes after LPS treatment followed a similar pattern, with minimal differences at lower LPS doses but a significant fold change increase after 10 µg/mL LPS of both *IL8* (3.4, p = 0.04) and *CTGF* (2.4, p = 0.05) (**Figure 4D)**.

**Fig. 4.**
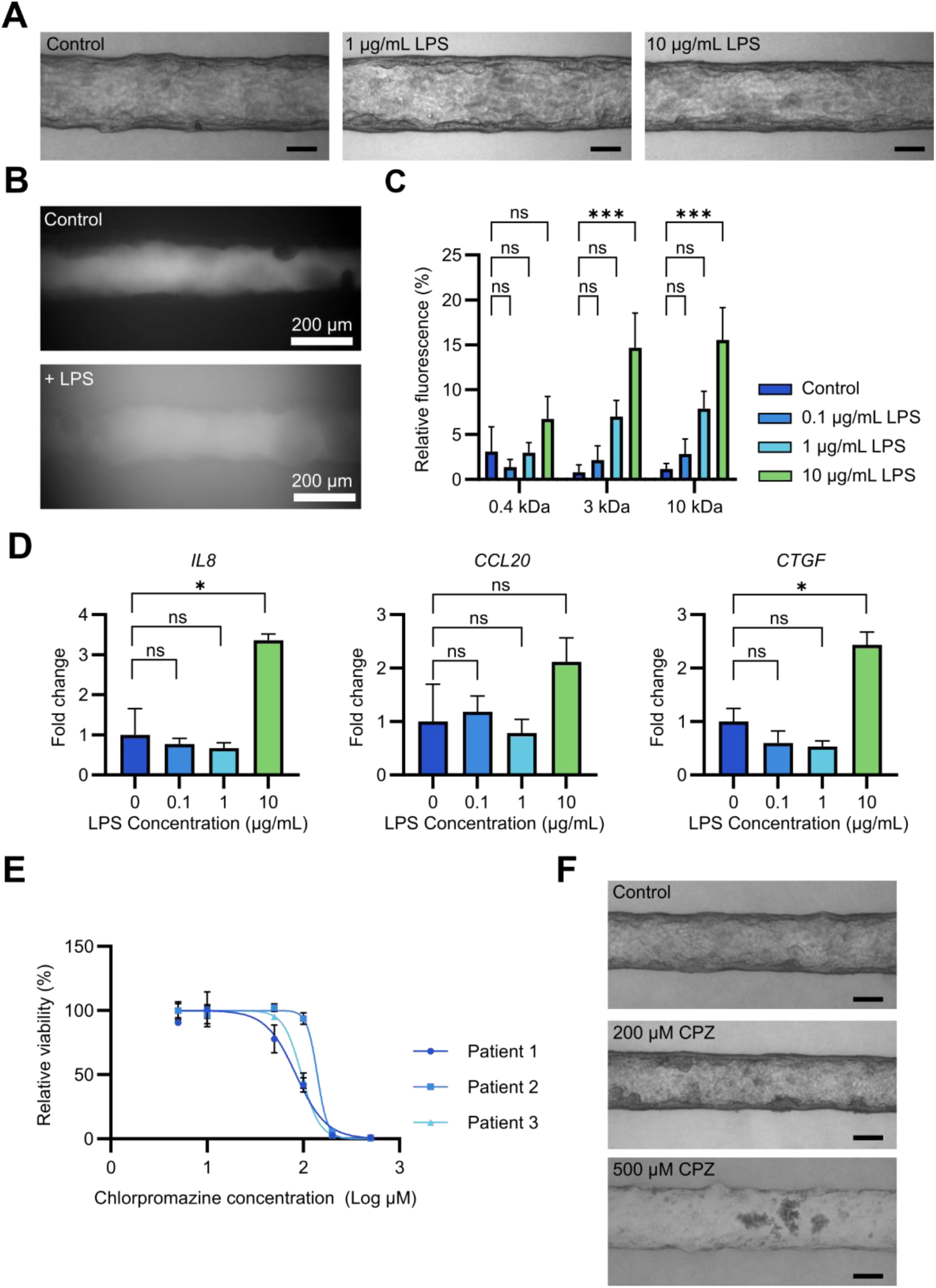
Response of the duct chip to injury. (A) Channels in the bile duct chip after treatment with LPS. (B) Confocal microscope images of 10 kDa dextran in channels with and without LPS 30 minutes after dextran addition. (C) Fluorescence in basal wells of different sized fluorescent compounds in channels after 24 hour treatment with different concentrations of LPS (Mean ± SEM). (D) Fold change of RNA for cytokines after treatment with different LPS concentrations measured with qPCR (Geometric mean ± SEM). (E) Viability of the channel measured with an MTS assay after treatment with different concentrations of chlorpromazine for 24 hours. (F) Channels after 24 hours of chlorpromazine treatment. All images and bar graphs are representative from one of three tested patient cholangiocyte lines (n = 3-4). Statistical analyses were calculated using one-way ANOVA. ns ≥ 0.05, ∗p < 0.05, ∗∗p < 0.01, and ∗∗∗p < 0.001, ∗∗∗∗p < 0.0001. CPZ, chlorpromazine; LPS, lipopolysaccharide.

### Chlorpromazine exposure can lead to drug induced liver injury in the chip

Drug induced liver injury (DILI) is a major problem in drug development and up to 40 % of DILIs involve cholestatic injury^25^. To assess whether our system could model drug induced bile duct damage we used CPZ, a drug known to cause damage to the liver and specifically the bile ducts *in vivo*^26, 27^.

Upon exposure to CPZ we observed dose-dependent toxicity in the bile ducts using an MTS assay (**Figure 4E**). Between individual patients, limited variability in toxicity was seen, with EC50 values of 84, 140 and 96 µM respectively for three different patients (**Table S6**). At a concentration of 200 µM CPZ damage to the epithelium and the presence of rounded dead cells can be observed with brightfield microscopy, and at 500 µM a complete loss of the bile duct is apparent (**Figure 4F)**.

### Treatment of biliatresone induced biliary atresia model

BA is an acute cholestatic disease presenting in neonates that has been associated with Biliatresone, a toxin known to induce BA in animals^28^. To test potential pharmacological treatment modalities for BA we challenged the chip with Biliatresone to induce treatable biliary changes.

After 48 hours of Biliatresone treatment, a range of morphological changes could be observed with increasing Biliatresone concentration. Numerous round cellular structures begin forming within the channels with a Biliatresone dose of 2 µg/mL and increasingly present at the higher dose of 5 µg/mL (**Figure 5A**). At 10 µg/mL a complete collapse of the channel is seen with detachment of the epithelium from the surrounding collagen gel (**Figure 5A**), morphologically resembling the collapse also seen in organoids at the same dose (**Figure S4A**). The round structures consist of dense groups of cells with disrupted actin polarization (‘Region 1’, **Figure 5B**), which aligns with previous studies demonstrating loss of actin polarization in some BA organoids^29^ and non-BA organoids after treatment with Biliatresone^30, 31^. Regions of epithelium with a reduced columnar phenotype were also frequently observed after Biliatresone treatment (‘Region 2’, **Figure 5B**). The Biliatresone phenotype could be partially ameliorated by treatment with NALC, an antioxidant and glutathione source which has shown clinical benefit in BA^32, 33^. When 100 µM NALC was added alongside Biliatresone challenge, increased viability and reduced number of internal structures was observed (**Figure 5C-E**) with a 160 % increase in viability measured for 2 µg/mL Biliatresone and 100 µM NALC (p < 0.0001), however the effect on overcoming the loss of viability after 5 µg/mL Biliatresone treatment was minimal (**Figure 5E**).

**Fig. 5.**
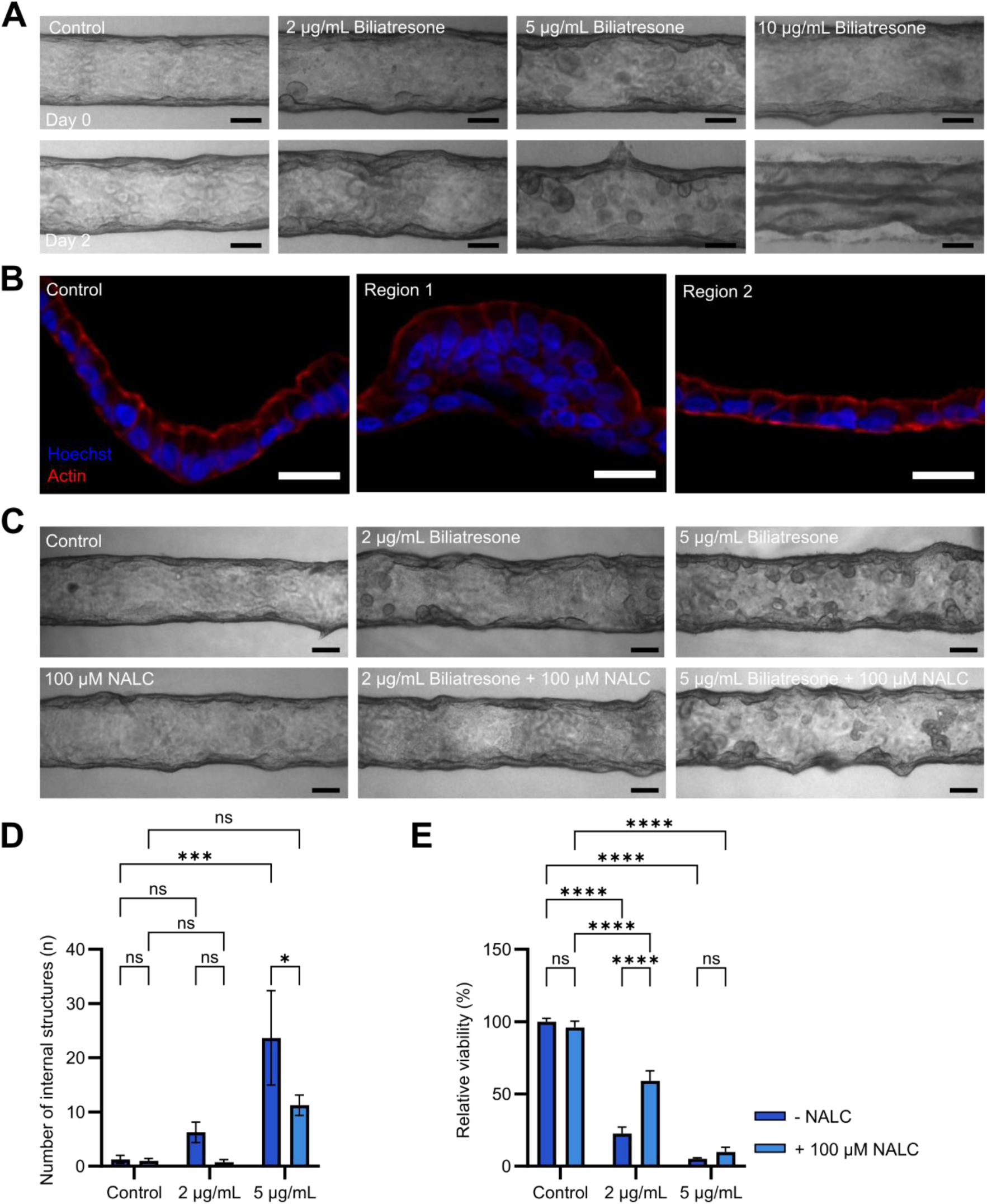
Treatment of the chip with biliatresone to induce a biliary atresia phenotype. (A) Bile duct channels before and after treatment with different concentrations of biliatresone. Scale bars are 100 µm. (B) Different morphological features present within biliatresone treated bile ducts. Scale bars are 20 µm. (C) Channels after treatment with different concentrations of biliatresone with or without 100 µM N-acetyl-L-cysteine. Scale bars are 100 µm. (D) Quantification of the number of internal structures within the bile duct channels after treatment. (Mean ± SEM). (E) Viability of the channel measured with an MTS assay after treatment with biliatresone and 100 µM N-acetyl-L-cysteine (Mean ± SEM). All images and graphs are representative data from one of 3 patient cholangiocyte lines tested (n = 4). Statistical analyses were calculated using one-way ANOVA. ns ≥ 0.05, ∗p < 0.05, ∗∗∗∗p < 0.0001. NALC, N-acetyl-L-cysteine.

## Discussion

Studies of bile duct diseases rely heavily on the use of mouse models that lack direct relevance to the human diseases^4^ and are therefore insufficient as tools to close the large unmet needs in cholangiopathies^1^.

Advanced *in vitro* techniques to replace or supplement mouse studies are increasingly being used, but also these approaches have drawbacks such as poor scalability and use of materials not suitable for drug studies limiting their applicability^15–17^. Establishment of an *in vitro* bile duct model allowing in-depth studies of biliary injury and disease at a suitable scale will dramatically increase the throughput and gain for future studies. In the present paper we demonstrate that we can produce an effective organ-on-a-chip model of the bile duct using patient-derived material which reliably mimics the structure and function of the bile duct. The translational potential of this system is demonstrated by investigating bile duct injury from various clinically relevant sources.

The cholangiocytes in our system form a functional epithelium with enhanced polarization compared to cholangiocyte organoids. Epithelial barrier function, a key aspect of bile duct function^34^, is effectively maintained over long time periods exhibiting a tight barrier to a range of different sized dextran molecules. The barrier function is further strengthened by mucus secretion on the apical surface of the cholangiocytes. Future studies could investigate inclusion of patient- and disease-matched bile and cholangiocytes to study the interplay between these two factors within diseases, something which is not achievable with animal models and classical *in vitro* assays.

Cholangiocytes *in vivo* are continuously exposed to microbial products in bile with low-grade colonization, which occurs in around 50% of PSC cases^35, 36^, and they constitutively express Toll-like receptors (TLRs) capable of sensing pathogen-associated signals^37, 38^. TLR activation has been implicated in several biliary diseases such as PBC and PSC^39, 40^, supporting a role for microbial signaling in biliary injury, and specific microbial species have been linked to poorer transplant-free survival in PSC^41, 42^. Despite these findings, mechanistic insights into how microbial signals affect cholangiocyte function, an interaction that a bile duct on a chip is ideal for studying, remain limited. As proof of concept for such studies, we added the gram-negative bacterial component LPS into the apical wells of the chip. When treated with LPS, a dose-dependent response was seen with leakage of dextran molecules increasing significantly at 10 µg/mL LPS, the highest dose tested, and non-significant variation seen at lower LPS concentrations. On the other hand, no significant change in the leakage of LY was seen with any LPS concentration. As LY is a smaller molecule and exhibits a higher baseline permeability across the epithelium, more significant disruption of the epithelial barrier may be required to observe changes with the level of sensitivity these assays allow. The production of cytokines was also increased after LPS treatment in the same pattern as for the barrier leakage, with 10 µg/mL LPS required to induce a significant effect. These data demonstrate that while LPS can have an effect on the barrier integrity of the bile duct *in vitro*, the barrier is very tolerant to LPS challenge with the levels of LPS needed to induce a significant response representing extreme levels found in the bile *in vivo*^43^.

Drug induced liver injury represents a major clinical challenge and is currently the leading cause of post-market withdrawal of drugs^44^. The major focus of studies of DILI has been on hepatocellular damage, but up to 40 % of DILI cases present with some degree of cholestatic injury^45^ highlighting that probing for bile duct related affection prior to clinical studies would be of significant value. Using CPZ, a drug which has been found to cause DILI including damage to the bile ducts^46^, we observed clear dose dependent cytotoxicity at comparable doses to those previously reported for cholangiocyte organoids^47^. In our model CPZ is dosed into the interior of the bile duct, a likely route of exposure for cholangiocytes *in vivo* due to the biliary secretion of drugs and their metabolites by hepatocytes. Use of our scalable system in this way opens up for testing a range of different drugs and their metabolites in parallel, which would be of relevance for the pharmaceutical industry as a significant proportion of drugs affecting cholangiocytes will have been metabolized by the hepatocytes prior to secretion into the bile ducts^44^. The use of patient derived cells also allows for a personalized medicine approach to patient treatment, with the testing of potential treatments on an individual basis having the potential to overcome some of the difficulties in treating highly heterogeneous diseases like cholangiopathies.

For disease modelling and treatment we focused on BA since the pathological phenotype could be explored using a cholangiocyte-only bile duct model and BA has been shown to be inducible with the toxin Biliatresone both *in vivo* and *in vitro*^28, 30, 48^. Addition of Biliatresone to our system led to the formation of organoid-like structures within the channels, along with a corresponding reduction in viability. While no major changes were seen to polarization as measured using staining for actin, expression of actin within the internal structures suggested poor polarization, and regions of the epithelia showed a less columnar morphology after Biliatresone treatment than the untreated epithelia. While not performed in this study, an future advantage of using the bile duct on a chip system for modelling this toxicity rather than organoids is the flexibility to deliver apical exposure to the cholangiocytes, the mode which has been hypothesized to be necessary for Biliatresone toxicity *in vivo*^49^. The addition of NALC, a compound known to reduce oxidative stress and provide a source of glutathione, led to a clearly improved morphology and reduced the loss of viability caused by a low dose of biliatresone but could not overcome the damage caused by a dose of 5 µg/mL. This finding strikingly aligns with results from clinical trials when NALC was tested^33, 50^. Collectively these data demonstrate that it is possible to induce human disease-relevant damage to the *in vitro* bile duct and even explore potential therapeutic treatment options. Future studies could further develop this disease model and introduce a range of different analytical measurements to allow for more in-depth analysis of the mechanisms and impact of treatment. While the data support the use of this system for modelling a range of pathological processes, there are still limitations which can be overcome to allow more complex studies to be performed. The system is lacking other key components of bile duct diseases, in particular the circulating immune system which precludes studies into immune-cholangiocyte interactions.

In this study we have for the first time demonstrated the feasibility of using an *in vitro* bile duct to model a range of disease mechanisms including bacterial contamination, drug toxicity and BA. The effects of these conditions can be readily measured in the chip and potential treatment options explored, with the reproducibility and scalability of the system allowing this to be done at a scale relevant for more complex studies. Future work would aim to look in more depth at the underlying mechanisms behind the conditions tested and test potential therapeutic treatments to cover a range of possible mechanisms, as well as between different patients with potential for a personalized medicine approach to treatment.

## Acknowledgements

Confocal imaging was performed at the Norwegian Centre for Stem Cell Research, Oslo University Hospital. We thank Oda Helgesen Ramberg for providing laboratory support, Marie Christin Röcklinger for support with organoid culture, Liv Wenche Thorbjørnsen and Sissel Åkra for coordinating collection of patient material and Merete Tysdahl for providing administrative support.

## Author Contributions

**Henry W. Hoyle:** Conceptualization, methodology, Investigation, Writing - Original Draft, Funding Acquisition. **Anna K. Frank:** Conceptualization, Methodology, Investigation, Funding Acquisition. **Enya Amundsen-Isaksen:** Methodology, Investigation. **Sarah Peisl:** Conceptualization, Methodology, Investigation. **Oline Ø. Hovland:** Methodology, Investigation. **Jeremy Yeoh:** Investigation. **Mughilan Selvarajah:** Investigation. **Aleksandra Aizenshtadt:** Conceptualization. **Kayoko Hirayama-Shoji:** Conceptualization, Methodology. **Fotios Sampaziotis:** Conceptualization. **Tom H. Karlsen:** Conceptualization, Resources. **Mathias Busek:** Conceptualization. **Stefan Krauss:** Conceptualization, Writing - Review & Editing, Resources. **Espen Melum:** Conceptualization, Methodology, Resources, Writing - Original Draft, Project Administration, Funding Acquisition.

## Data Transparency Statement

Additional data, methods and consumables information are available in the supplementary information. Full datasets are available upon reasonable request to the corresponding author.

## Abbreviations

AS: Alagille syndrome
BA: Biliary Atresia
BSA: Bovine serum albumin
CPZ: Chlorpromazine
CLF: Cholyl-lys-fluorescein
DILI: Drug induced liver injury
DKK1: Dickkopf-related protein 1
EPCAM: Epithelial cell adhesion molecule
ERCP: Endoscopic retrograde cholangiopancreatography
FBS: Fetal bovine serum
hEGF: human epidermal growth factor
LPS: Lipopolysaccharide
LY: Lucifer yellow
MTS: [3-(4,5-dimethylthiazol-2-yl)-5-(3-carboxymethoxyphenyl)-2-(4-sulfophenyl)-2H- tetrazolium
NALC: N-acetyl-L-cysteine
OoC: Organ on a chip
PAMP: Pathogen-associated molecular pattern
PBC: Primary biliary cholangitis
PBS: Phosphate buffered saline
PDMS: polydimethylsiloxane
PFA: Paraformaldehyde
PLL: Poly-L-lysine
PS: Polystyrene
PSC: Primary sclerosing cholangitis
RT-qPCR: Reverse transcription quantitative polymerase chain reaction
SNA: Sambucus Nigra
TLR: Toll-like receptor
ZO-1: Zonula Occludens-1

