## Supplemental File 1 for "Modelling mechanisms and treatment of cholangiopathies with a bile duct on a chip"

### Supplementary Materials and Methods

*Patient Material*

Cholangiocytes were obtained from the bile ducts of patients undergoing ERCP. The patients were recruited at the Section of Gastroenterology at the Department of Transplantation Medicine, Oslo University Hospital Rikshospitalet. Patient inclusion in the study was performed independent of gender (**Table S1**).

Written informed consent was obtained prior to ERCP and the collection and use of material by the Norwegian PSC research center biobanks has been approval by the Regional Ethical Committee (REK 15368, 599798 and 18221).

*Culture of ERCP-derived cholangiocyte organoids*

Cholangiocyte organoids were derived from ERCP brushes as previously described [1]. All organoids were derived from patients diagnosed with PSC.

Briefly, brush cytology samples from ERCP procedures were transferred into Williams medium E (Gibco, Thermo Fisher Scientific, Waltham, MA, USA) supplemented with 10 mM nicotinamide (Sigma-Aldrich, St Louis, MO, USA), 17 mM sodium bicarbonate (Sigma-Aldrich), 0.2 mM 2-Phospho-L-Ascorbic Acid Trisodium Salt (Gibco), 14 mM D-glucose (Sigma-Aldrich), 0.1 µM dexamethasone (Gibco), 2,12 mM L-glutamine (Gibco), 6.3 mM sodium pyruvate (Gibco), 20 mM HEPES (Invitrogen, Thermo Fisher Scientific), 1X Insulin-Transferrin-Selenium (Gibco), 50 U/mL penicillin and 50 µg/mL streptomycin (Gibco) (Wash medium) supplemented with 50 ng/mL human epidermal growth factor (hEGF, Bio-Techne Minneapolis, MN, USA) and 10 µM Y-27632 (TargetMol Chemicals Inc., Boston, MA, USA). Cells were removed from brushes by pipetting and the cell suspension centrifuged for 4 minutes at 482 g to pellet cells. After removal of supernatant, cells were resuspended in wash medium with 1500 ng/mL R-spondin 1 (PeproTech, Thermo Fisher Scientific), 300 ng/mL Dickkopf-related protein

1 (DKK1, MedChemExpress, Monmouth Junction, NJ, USA), 30 µM Y-27632, 150 ng/mL hEGF, and 6 µM forskolin (Sigma-Aldrich). Suspended cells were mixed 1:2 with phenol red-free Matrigel matrix (Corning Inc., New York, USA) and pipetted into wells of a 24-well plate. Plates were incubated upside-down at 37 °C for 30 minutes to allow Matrigel to polymerize before addition of wash medium supplemented with 500 ng/mL R-spondin 1, 100 ng/mL DKK1 and 50 ng/mL hEGF (Growth medium) along with 10 µM Y-27632, and 2 µM forskolin. All samples were routinely tested for mycoplasma (MycoAlert PLUS, Lonza, Basel, Switzerland) to ensure no contaminations were present.

*Fixation and sectioning*

Medium was aspirated from all wells, the chips were washed once with PBS and fixed with 4% formaldehyde (Thermo Fisher) at 4 °C for 20 minutes. Samples were subsequently washed in PBS three times for 5 minutes each. The coverslips were removed from the base of the chips and collagen gel removed and placed into optimum cutting temperature compound in a plastic embedding mold. Samples were frozen and stored at -80 °C.

Samples were sectioned at 10 µm thickness using a Leica CM3050 S cryostat (Leica Biosystems, Nussloch, Germany) onto Superfrost™ Plus microscope slides (Thermo Fisher). Sections were stored at -20 °C until staining was performed.

*Immunofluorescent staining*

Blocking was performed for 1 hour with 5 % bovine serum albumin (BSA, Sigma-Aldrich) and 0.1% Triton X-100 (VWR, Radnor, PA, USA) in PBS. After blocking, primary antibodies were added at required dilutions in PBS containing 1 % BSA and 0.1 % Tween-20 (Bio-Rad, Hercules, CA, USA) (**Table S2**). Samples were incubated overnight at 4 °C. Sections were washed in PBS 3 times for 5 minutes each and secondary antibodies (**Table S3**) were added at a concentration of 1:1000 in PBS containing 1 % BSA and 0.1 % Tween-20. Samples were incubated for 2 hours at room temperature in the dark. Three further 5-minute washes in PBS were performed. For samples with actin or sambucus nigra (SNA) lectin staining, 0.1 % CellMask™ Deep Red actin tracking stain (Invitrogen) or 20 μg/mL fluorescein-tagged SNA lectin (Invitrogen) in PBS was added to the samples and incubated at room temperature for 15 minutes. Samples were washed twice for 5 minutes with PBS. #1.5 Glass coverslips were mounted to the slides using ProLong™ Glass antifade mountant with NucBlue™ (Invitrogen). Samples were stored at 4 °C prior to imaging. Imaging was performed using a Zeiss LSM 710 confocal microscope (Zeiss).

*Barrier permeability assay*

The epithelial barrier was tested by addition of Lucifer Yellow (LY, Invitrogen) at 100 μg/mL alongside 3 kDa Cascade Blue dextran and 10 kDa fluorescein dextran (Invitrogen) at concentrations of 500 μg/mL each. Medium was aspirated from the chips and 150 μL of fresh growth medium was added to the basal wells. 150 μL of fluorescent compound-containing medium was added to each apical well. Samples were incubated at 37 °C for one hour. Medium was removed from the basal reservoirs and the fluorescent intensity measured using a BioTek H1 plate reader at the required wavelengths for each fluorophore (**Table S4**)

*Rhodamine 123 transport assay*

Medium was aspirated from samples and fresh growth medium added to the apical wells. 5 μM Rhodamine 123 in growth medium was added to the basal wells and samples were incubated for 30 minutes. Medium from the apical wells was then removed and the fluorescent intensity measured at the necessary wavelengths using a BioTek H1 plate reader (**Table S4**). For samples to be imaged, fresh growth medium was added to the basal wells and samples incubated for a further 40 minutes at 37 °C before imaging on a Zeiss LSM 710 confocal microscope.

For Verapamil treatment, prior to addition of rhodamine 123 the samples were treated with growth medium containing 20 µM of Verapamil (Sigma-Aldrich). Samples were incubated with verapamil for 30 minutes at 37 °C. Control samples were treated the same but with fresh growth medium not containing verapamil. After incubation, samples were washed with growth medium and the rhodamine 123 assay performed.

*Cholyl-lys-fluorescein transport assay*

Samples were incubated for 30 minutes with wash medium alone or containing 50 µM Linerixibat (MedChemExpress). Samples were washed and 100 µL of wash medium was added to the basal wells and 100 µL wash medium with 10 µM cholyl-lys-fluorescein (CLF, Avanti Research, Alabaster, AL, USA) was added to the apical wells. Samples were incubated for 30 minutes at 37 °C before sampling the medium in the basal wells and reading the fluorescence at the necessary wavelengths using a BioTek H1 plate reader (**Table S4**). For imaging, the apical wells were washed with fresh wash medium and the samples imaged using a Zeiss LSM 710 confocal microscope.

*Chlorpromazine treatment*

Culture medium was aspirated from the chip and fresh growth medium containing CPZ (Sigma-Aldrich) at the desired dose, ranging from 1 µM to 500 µM was added to the apical wells, while fresh growth medium without CPZ was added to the basal wells. Chips were incubated for 24 hours before imaging with a brightfield microscope and running viability assays.

*Lipopolysaccharide treatment*

Culture medium was aspirated and 150 µL of growth medium containing different concentrations (0.1, 1,10 µg/mL) of LPS (Invitrogen) added to the apical wells. Fresh growth medium was added to the basal wells. Samples were incubated for 24 hours before analysis by brightfield microscopy and reverse transcription quantitative polymerase chain reaction (RT-qPCR) was performed.

*Biliatresone treatment*

Fresh growth medium containing BSO (MedChemExpress) at concentrations from 2 µg/mL to 10 µg/mL was added to apical and basal wells. For additional N-acetyl-L-cysteine (NALC, Thermo Fisher) treatment, NALC was also added at 100 µM to the medium simultaneously with BSO. Samples were incubated for 48 hours before analysis was performed.

*Cell viability assay*

The viability of cells was measured using the CellTiter 96® assay based on the reduction of [3-(4,5-dimethylthiazol-2-yl)-5-(3-carboxymethoxyphenyl)-2-(4-sulfophenyl)-2H-tetrazolium (MTS)(Promega, Madison, Wisconsin, USA). A solution of 1 mg/mL MTS and 0.1 mg/mL phenazine ethosulfate (PES)(Sigma-Aldrich) was made up in wash medium. Culture medium was aspirated from plates and the MTS solution added to each well. For the chip, reagent was added both to the apical and basal wells. Samples were incubated at 37 °C for 1 hour. The supernatant was briefly mixed and diluted 1:5 into PBS in a 96-well plate in duplicate. The absorbance was read at 490 nm using a Synergy H1 plate reader.

*RT-qPCR*

RNA isolation was performed using a Reliaprep™ RNA tissue miniprep system (Promega). Medium was aspirated from the chips and 150 µL of ice cold LBA + TGA buffer was added to the channels. The collagen gel was disrupted with a 21 G needle and transferred to a 2 mL tube. The remainder of the protocol was carried out following the manufacturer's protocol. RNA quality and quantity was measured using a DS-11

Spectrophotometer (DeNovix, Wilmington, DE, USA) and a Qubit 4 Fluorometer (Invitrogen) with a Qubit RNA High sensitivity assay kit. Primers used are as listed in Table S5 (**Table S5**).

cDNA synthesis was performed using a high capacity cDNA reverse transcription kit (Thermofisher)following the manufacturers protocol with a reaction volume of 20 µL.

SYBR Green based qPCR was performed using GoTaq 2x qPCR Master Mix (Promega) following the manufacturers protocol. All samples were diluted to 0.25 ng/µL cDNA and a reaction volume of 15 µL was used with 2 replicates per sample. Genes were normalized to the housekeeping gene *GAPDH* followed by the mean of the organoid samples to give a ΔΔCT value. Fold change values were calculated with the formula

$$\text{fold change} = 2^{-\Delta\Delta CT}.$$

Values are presented as geometric mean ± geometric standard error.

*Image analysis*

Brightfield images were captured using a Nikon Eclipse TS100 microscope (Nikon, Tokyo, Japan) with a Teledyne Moment camera (Teledyne Vision Solutions, Ontario, Canada). Images were blinded and the number of internal structures measured in ImageJ 1.53t [2]. Criteria for internal structures were rounded structures within the focal plane which have over half the structure separate from the epithelium.

Cell height and width were calculated from samples stained for F-actin to outline the cells as described in the immunofluorescent staining section. Images were taken with a Zeiss LSM 710 confocal microscope and measurements were taken manually on blinded images using ImageJ 1.53t. 20 cells were measured per sample, with three samples measured per condition, and repeated for each patient line.

*Flow rate calculation*

The fluid flow rate in the chip was calculated using a modified version of the Hagen-Poiseuille equation assuming that the low flow rate through the channel is laminar. The equation used to calculate the flow velocity, $v$, over time is

$$v = \frac{R^2 \rho g \sin(\theta)}{8\mu}$$

Where $R$ is the radius of the channel, $\rho$ is the density of the culture medium, $g$ is the acceleration due to gravity, $\theta$ is the angle of the rocking platform and $\mu$ is the dynamic viscosity of the culture medium. The angle, $\theta$, was modelled over time, $t$, using a triangle wave equation

$$\theta(t) = \frac{4a}{p}\left|\left(\left(t - \frac{p}{4}\right) \bmod p\right) - \frac{p}{2}\right| - a$$

where $a$ is the amplitude of the wave and $p$ is the period. Upper and lower limits of 12 degrees were used to define the 10 second pause between changing directions.

*Drug absorption assay*

To test the absorption of compounds to different cell culture materials, deoxycholic acid-iFluor 647 conjugate (DCA-647, AAT Bioquest, Pleasanton, CA, USA) was used due to its clinical relevance, hydrophobicity and fluorescence outside of the autofluorescent range of the 3D printed resin. Polystyrene cell culture plates were used as a control. For PDMS, Sylgard 184 (Dow Inc., Midland, MI, USA) was mixed following the manufacturer’s instructions, 20 µL added per well to a 96-well plate, followed by curing at 60 °C for 4 hours. For the resin, 20 µL of Formlabs High Temperature Resin V2 was added per well to a 96-well plate and cured in a FormCure for 20 minutes at 80 °C. Plates were washed with isopropanol prior to use. 100 µL of 50 µM DCA-647 was added per well and incubated at 37 °C for 24 hours. Wells were washed 5 times with PBS for 10 minutes each before 100 µL of fresh PBS being added and the fluorescence measured at the necessary wavelengths (**Table S4**).

*Statistics*

All experiments were performed with cholangiocytes from 3 different PSC patients, with 3 to 4 channels for each condition unless otherwise stated. Data is presented as mean ± SEM unless otherwise stated. Statistics were calculated using GraphPad Prism version 10.4.1 (GraphPad, San Diego, CA, USA). Comparisons between two groups were performed with Welch's t-test while comparisons between multiple groups were performed using one-way ANOVA with Dunnett's test for post-hoc comparison of treatment to control groups. Comparisons within grouped analyses were performed with two-way ANOVA using Šídák's multiple comparisons test for post hoc comparisons between group means.

### Supplementary Figures

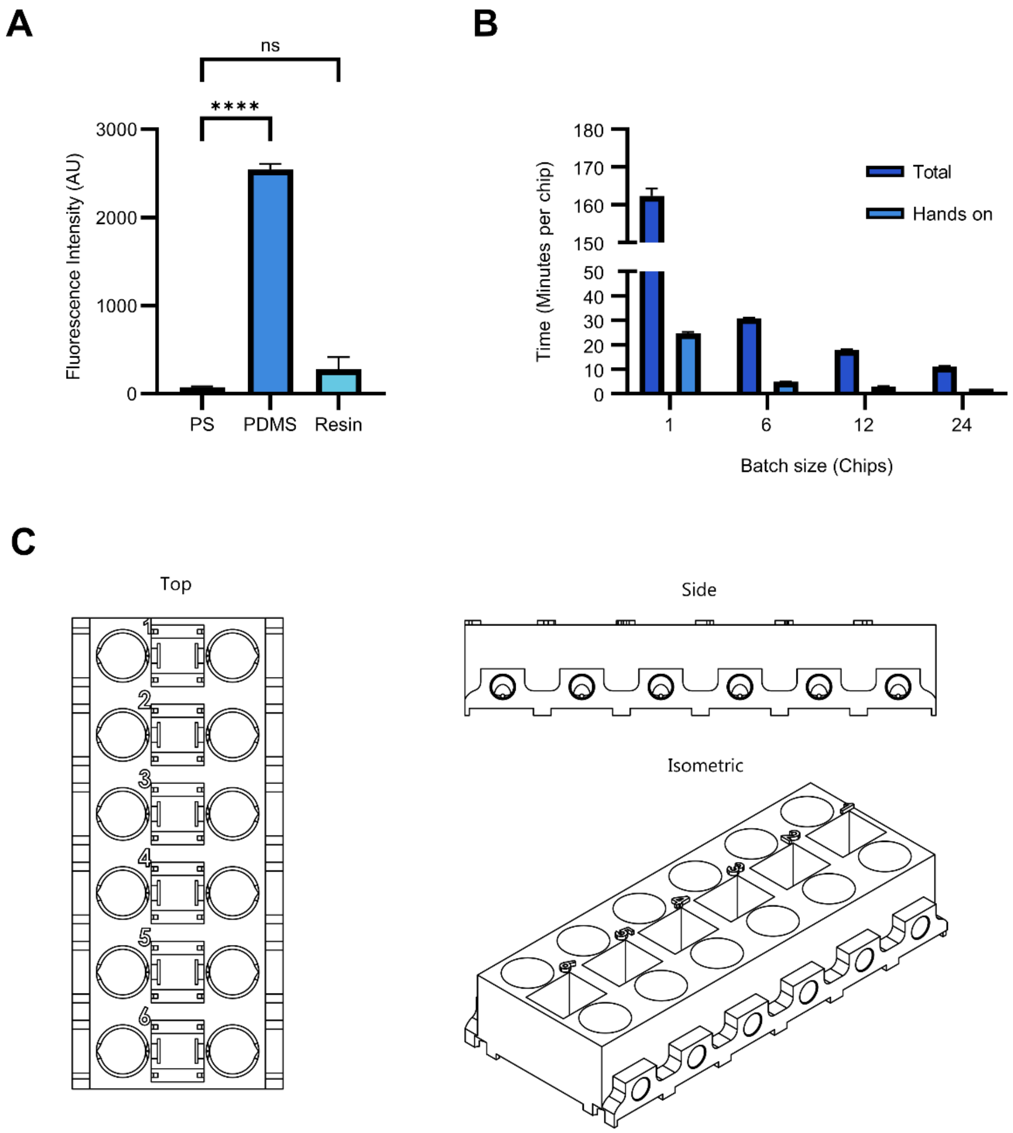

**Fig. S1. Design properties of the bile duct chip.**

(A) Absorption of deoxycholic acid-iFluor 647 conjugate to different plastics used for cell culture (Mean ± SEM, n = 5). (B) Time taken to fabricate chips compared to the batch size. (Mean ± SD, n = 3) (C) Schematics of the bile duct chip. Statistical analysis was calculated using one-way ANOVA. ns ≥ 0.05, ****p < 0.0001. PDMS, Polydimethylsiloxane; PS, Polystyrene.

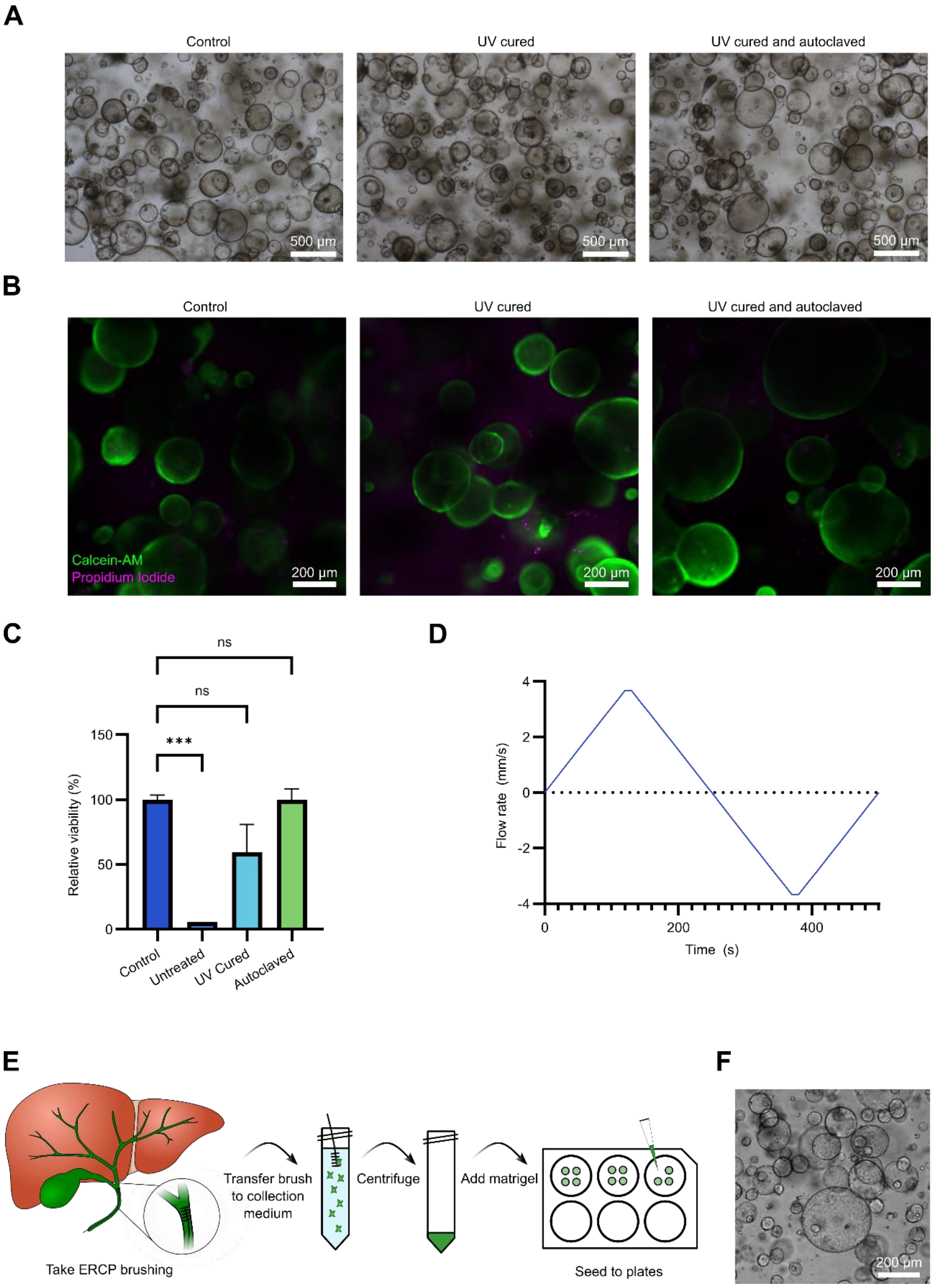

**Fig. S2. Biocompatibility of 3D printer resin with cholangiocytes**

(A) Representative brightfield images of cholangiocyte organoids after exposure to 3D printed plastic with different treatments. (B) Representative confocal microscope images of live and dead cell staining of cholangiocyte organoids using calcein-AM and propidium iodide staining after exposure to 3D printed plastic with different treatments. (C) Viability of the cholangiocyte organoids measured with an MTS assay after exposure to 3D printed plastic with different treatments (Mean ± SEM, representative data from one of three patient cholangiocyte lines, n = 4). (D) An estimate of the flow rate through the bile duct channel as a function of time calculated using a modified form of the Hagen–Poiseuille equation. (E) Schematic protocol for the derivation of cholangiocyte organoids from endoscopic retrograde cholangiopancreatography. (F) Representative brightfield microscope image of cholangiocyte organoids. Statistical analysis was calculated using one-way ANOVA. ns ≥ 0.05, $****p < 0.0001$. ERCP, endoscopic retrograde cholangiopancreatography; UV, ultraviolet.

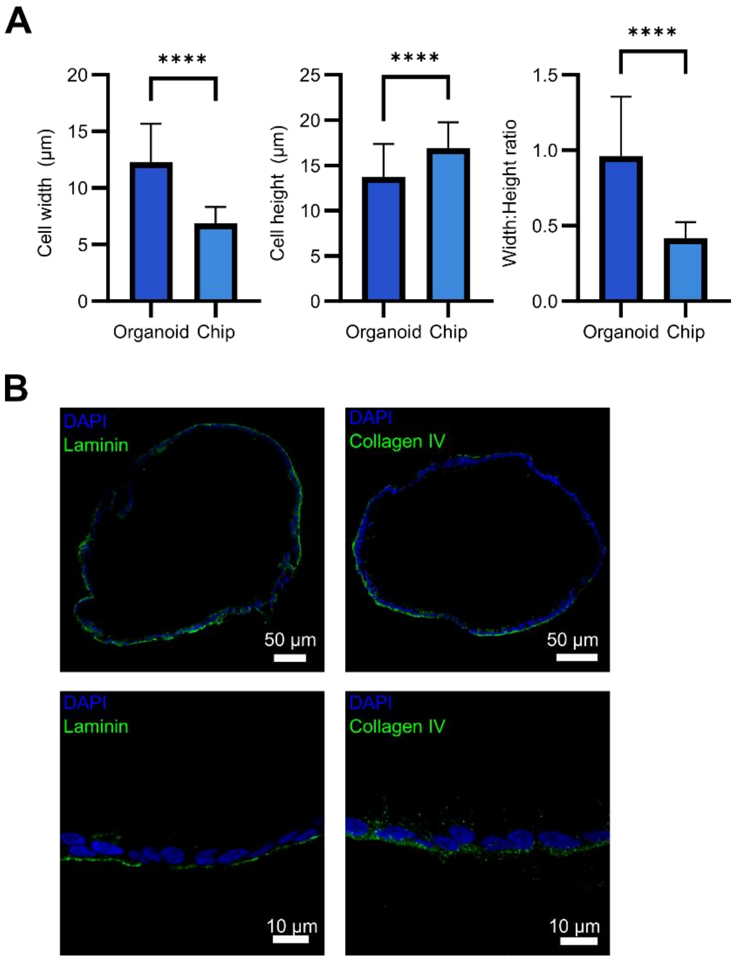

**Fig. S3. Characterization of organoids in the duct chip**

(A) Quantification of cholangiocyte dimensions when cultured as organoids compared to in the chip. (Mean ± SD, data pooled from 3 different patient lines, each n = 4) (B) Representative confocal microscope images from one of 3 tested patient cholangiocyte lines (n=4) of immunofluorescently stained basement membrane proteins. Proteins of interest are labelled in green and nuclear Hoechst 33342 staining is labelled in blue. Statistical analyses were calculated using Welch's t test. ns ≥ 0.05, ****p < 0.0001.

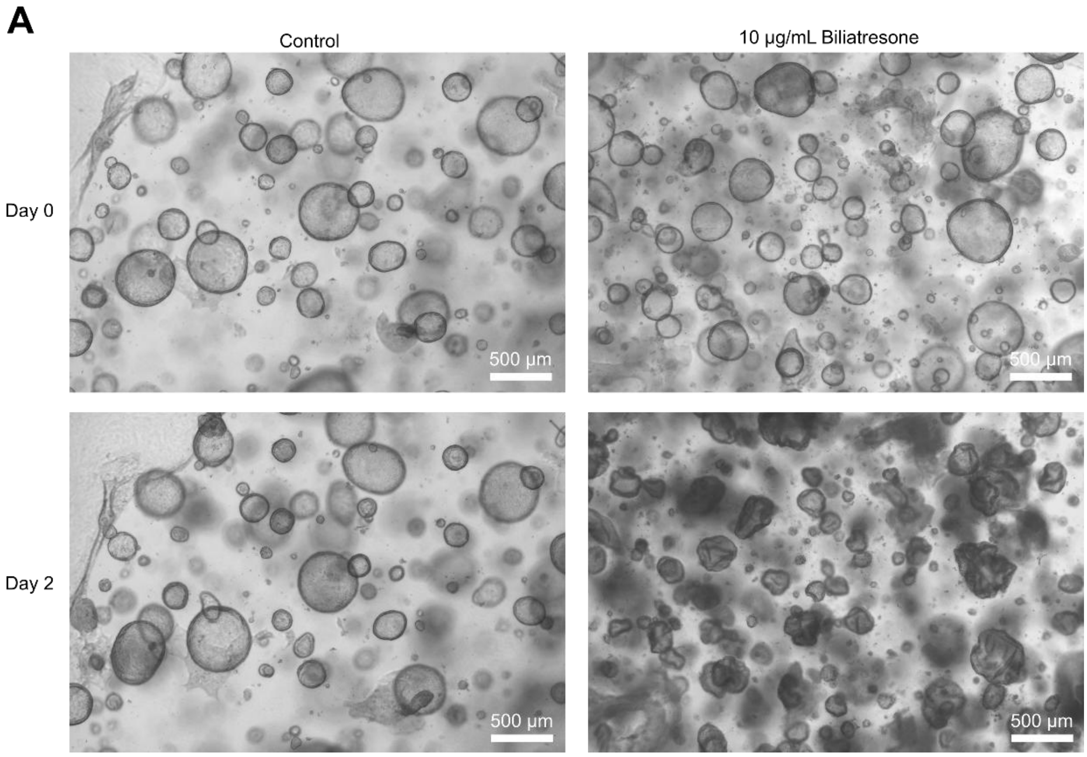

**Fig. S4. Biliatresone treatment of cholangiocyte cultures**

(A) Representative brightfield microscope images cholangiocyte organoids treated with 10 µg/mL biliatresone for 48 hours. Images are representative from one of three tested patient cholangiocyte lines.

### Supplementary Tables

#### Table S1: Patient details

| Patient # | Diagnosis | Age (Years) | Sex | IBD |
|---|---|---|---|---|
| 1 | PSC | 49 | Female | None |
| 2 | PSC | 56 | Male | UC |
| 3 | PSC | 50 | Male | UC |
| 4 | PSC | 57 | Male | UC |
| 5 | PSC | 36 | Male | None |

**Table S2: Primary Antibodies**

| Antibody | Supplier | Catalog # | Dilution | Host species |
|---|---|---|---|---|
| Claudin 2 | Invitrogen | 51-6100 | 1:100 | Rabbit |
| Collagen IV | Abcam | ab6586 | 1:200 | Rabbit |
| Cytokeratin 19 | Abcam | ab52625 | 1:200 | Rabbit |
| Cytokeratin 7 | Abcam | ab181598 | 1:200 | Rabbit |
| E-Cadherin | Abcam | ab231303 | 1:100 | Mouse |
| EPCAM | Abcam | ab71916 | 1:200 | Rabbit |
| Laminin | Abcam | ab11575 | 1:200 | Rabbit |
| ZO-1 | Abcam | ab221547 | 1:200 | Rabbit |

**Table S3: Secondary Antibodies**

| Target species | Supplier | Catalog # | Dilution | Host species | Fluorophore |
|---|---|---|---|---|---|
| Rabbit | Invitrogen | A10037 | 1:1000 | Donkey | Alexafluor Plus 488 |
| Mouse | Invitrogen | A-21206 | 1:1000 | Donkey | Alexafluor 568 |

**Table S4: Plate reader wavelengths**

| Fluorophore | Catalog # | Supplier | Excitation | Emission |
|---|---|---|---|---|
| Lucifer Yellow | L0144 | Sigma-Aldrich | 430 | 545 |
| 3 kDa Cascade Blue dextran | D7132 | Invitrogen | 400 | 430 |
| 10 kDa fluorescein dextran | D1821 | Invitrogen | 500 | 530 |
| Cholyl-lys-fluorescein | 810389P | Avanti Research | 485 | 528 |
| Rhodamine 123 | 83702 | Sigma-Aldrich | 485 | 528 |
| Deoxycholic acid-iFluor 647 | 36705 | AAT Bioquest | 645 | 670 |

**Table S5: RT-qPCR Primers**

| Gene | Forward | Reverse |
|---|---|---|
| AE2 | TCCAAGCACGAGCTGAAACTG | CACTGCGTCCAACTCCACA |
| AQP1 | TAACCCTGCTCGGTCCTTTG | AGTCGTAGATGAGTACAGCCAG |
| ASBT | GGACAATGCAACAGTTTGCTC | CCGTACTTAGGACCACACTTAGG |
| CFTR | CCTATGACCCGGATAACAAGGA | GAACACGGCTTGACAGCTTTA |
| CLDN2 | GCCTCTGGATGGAATGTGCC | GCTACCGCCACTCTGTCTTTG |
| CLDN3 | AACACCATTATCCGGGACTTCT | GCGGAGTAGACGACCTTGG |
| GAPDH | ACAACTTTGGTATCGTGGAAGG | GCCATCACGCCACAGTTTC |
| MDR1 | GGGAGCTTAACACCCGACTTA | GCCAAAATCACAAGGGTTAGCTT |
| MUC1 | TGCCGCCGAAAGAACTACG | TGGGGTACTCGCTCATAGGAT |
| TJP1 | ACCAGTAAGTCGTCCTGATCC | TCGGCCAAATCTTCTCACTCC |

**Table S6: EC50 values**

| Patient # | EC50 |
|---|---|
| 1 | 84.1 ± 1.1 |
| 2 | 140 ± 1.1 |
| 3 | 96.0 ± 1.0 |
